# Primary cilia shape postnatal astrocyte development through Sonic Hedgehog signaling

**DOI:** 10.1101/2024.10.17.618851

**Authors:** Rachel Bear, Steven A. Sloan, Tamara Caspary

**Author notes:** Address correspondence to Tamara Caspary.

## Abstract

Primary cilia function as specialized signaling centers that regulate many cellular processes including neuron and glia development. Astrocytes possess cilia, but the function of cilia in astrocyte development remains largely unexplored. Critically, dysfunction of either astrocytes or cilia contributes to molecular changes observed in neurodevelopmental disorders. Here, we show that a sub-population of developing astrocytes in the prefrontal cortex are ciliated. This population corresponds to proliferating astrocytes and largely expresses the ciliary protein ARL13B. Genetic ablation of astrocyte cilia *in vivo* at two distinct stages of astrocyte development results in changes to Sonic Hedgehog (Shh) transcriptional targets. We show that Shh activity is decreased in immature and mature astrocytes upon loss of cilia. Furthermore, loss of cilia in immature astrocytes results in decreased astrocyte proliferation and loss of cilia in mature astrocytes causes enlarged astrocyte morphology. Together, these results indicate that astrocytes require cilia for Shh signaling throughout development and uncover functions for astrocyte cilia in regulating astrocyte proliferation and maturation. This expands our fundamental knowledge of astrocyte development and cilia function to advance our understanding of neurodevelopmental disorders.

## INTRODUCTION

Astrocytes play critical roles in coordinating neural circuit development by communicating with neurons and other glia (Akinlaja & Nishiyama, 2024). Astrocytes are actively involved in many processes including synaptogenesis, synapse pruning, modulating neurotransmission, and regulating synaptic plasticity (Allen & Eroglu, 2017). Our knowledge of the complex functions that astrocytes serve in the developing brain continues to grow, yet our mechanistic understanding of the signals that guide how astrocytes, themselves, develop remains incomplete.

In rodents, cortical astrocytes are specified by radial glia progenitors during embryonic development (Ge et al., 2012). After birth, the astrocyte population keeps expanding through a combination of continued differentiation of radial glia and by local proliferation of committed astrocytes. (Clavreul et al., 2019). Astrocytes then begin a period of maturation, which includes changes in gene expression and morphology that is necessary for their functions at neuronal synapses (Bushong et al., 2004; Freeman, 2010; Zhang et al., 2016). Astrocytes acquire specialized structure and function across and within different brain regions resulting in considerable astrocyte heterogeneity (Batiuk et al., 2020; Bayraktar et al., 2020; Chai et al., 2017). Local environmental signals contribute to the differences among astrocytes (Farhy-Tselnicker et al., 2021; Lanjakornsiripan, 2018; Morel et al., 2014). These processes are crucial for astrocyte function such that disruptions to astrocyte development contribute to the molecular and morphological changes observed in neurodevelopmental disorders (Molofsky et al., 2012; Sloan & Barres, 2014). Therefore, identifying the molecular signals that underly astrocyte development is critical to our understanding of nervous system development and dysfunction.

Primary cilia are specialized signaling organelles that project from the cell membrane into the extracellular environment and are present on most cells (Pazour & Witman, 2003; S. P. Sorokin, 1968). In the brain, primary cilia are found on radial glia, neurons, astrocytes, and oligodendrocyte precursor cells (Bear & Caspary, 2024). The structure of cilia is well-suited for their signaling capabilities. Cilia are comprised of a microtubule-based axoneme that is assembled from the basal body by intraflagellar transport (IFT), the ciliary trafficking machinery (Klena & Pigino, 2022; Kozminski et al., 1993; Pazour et al., 2000; Rosenbaum & Witman, 2002). The axoneme is enclosed by the ciliary membrane and ciliary entry/exit is regulated by a complex of proteins that make up the transition zone at the base of the axoneme (Park & Leroux, 2022). The ciliary membrane localizes many different signaling receptors to coordinate rapid and efficient signal transduction within the cell (Anvarian et al., 2019; Christensen et al., 2012; Garcia-Gonzalo et al., 2015; Mykytyn & Askwith, 2017).

In mice, loss of cilia is embryonic lethal, highlighting the importance of cilia during embryonic development (Murcia et al., 2000). Mutations in ciliary genes lead to a class of human genetic disorders called ciliopathies, which present with a wide spectrum of symptoms that are often shared across multiple disorders (Reiter & Leroux, 2017; Turan et al., 2023). This includes neurological phenotypes such as neuroanatomical abnormalities, intellectual disability, epilepsy, and ataxia (Valente et al., 2014). However, the molecular mechanisms underlying these neurological impairments remain poorly understood and it is unclear whether astrocyte cilia contribute to these phenotypes.

Cilia modulate many different signaling pathways, including Sonic Hedgehog (Shh) signaling, that are critical for neuron development (Corbit et al., 2008; Ezratty et al., 2011; Huangfu et al., 2003; Lancaster et al., 2011; Schneider et al., 2005). Shh transduction requires cilia because the pathway components are dynamically trafficked within cilia to carry out their functions (Bangs & Anderson, 2017; Huangfu et al., 2003; Huangfu & Anderson, 2005; Liu et al., 2005; Rohatgi et al., 2007). Cilia were linked to neuron development upon the discovery that IFT proteins are required for Shh signaling, which patterns the neural tube (Echelard, 1993; Huangfu et al., 2003). This finding helped connect cilia to neuron proliferation because Shh signaling serves as a proliferative cue in cerebellar granule neurons (Chizhikov et al., 2007; Wechsler-Reya & Scott, 1999). The relationship between cilia and proliferation is supported by the dual function of centrosomes as the basal body as well as the microtubule organizing center. This explains the dynamic disassembly of cilia before cell division and reassembly of cilia in G_0_/G_1_ (Archer & Wheatley, 1971; S. Sorokin, 1962). In a distinct population of neurons, cilia function is necessary for interneuron migration, which highlights the diverse roles of cilia (Higginbotham et al., 2012). Critically, studies of cilia function in the brain have thus far focused on neurons and largely ignored their role in astrocytes.

To investigate potential signaling mechanisms involved in astrocyte development, we focused on primary cilia. Astrocyte primary cilia are found throughout the adult brain and vary in length and frequency by brain region (Kasahara et al., 2014; Ott et al., 2023; Sipos et al., 2018; Sterpka & Chen, 2018). Abnormal cilia structure is reported in a number of neurological diseases and brain injuries, but whether astrocyte function is impacted and how this affects disease pathology remains unknown (Ignatenko et al., 2023; Ma et al., 2022; Sterpka et al., 2020). Recent work has started to address the question of astrocyte function. Genetic disruption of astrocyte cilia alters the transcriptional signatures of cortical astrocytes in the adult brain (Wang et al., 2024). This results in disrupted actin dynamics, G-protein coupled receptor (GPCR) activity, and neuronal circuitry, ultimately leading to behavioral deficits. Additionally, genetic disruption of Shh signaling in astrocytes alters their expression of genes related to synapse regulation (Wang et al., 2024). Shh signaling is also known to act non-cell autonomously in astrocytes to regulate synapses and activity in specific populations of neurons (Hill et al., 2019; Xie et al., 2022). Layer V neurons are thought to be the main source of Shh ligand in the cortex, which may explain specific networks of neuron-astrocyte communication (Garcia et al., 2010; Harwell et al., 2012). Still, little is known about the role of cilia in developing astrocytes. This motivates our work to explore whether cilia function during astrocyte development and what signaling pathways they regulate in astrocytes.

To uncover the function of cilia during astrocyte development we generated two distinct *in vivo* models. We created genetic models using an inducible Cre-lox system to ablate astrocyte cilia as well as label astrocytes and their cilia. The first model targets immature astrocytes by inducing cilia ablation during embryonic development. The second model targets mature astrocytes by inducing cilia ablation during postnatal development. These independent cilia ablation models allow us to first investigate the role of astrocyte cilia in early stages of development and then specifically examine the function of cilia during astrocyte maturation. In this study, we identify that astrocyte cilia mediate Shh activity from early postnatal astrocyte development through maturation. We show that Shh activity is decreased in both immature and mature astrocytes upon cilia ablation. Finally, we determine the functional consequence of cilia ablation and find that cilia modulate astrocyte proliferation and morphology. This work reveals previously unknown roles for Shh signaling and cilia function during astrocyte development.

## METHODS

### Mouse lines

All mice were cared fwor in accordance with NIH guidelines and Emory University’s Institutional Animal Care and Use Committee (IACUC). Lines used were *Aldh1l1-Cre^ERT2^* [MGI:5806568, IMSR_JAX:031008] (Srinivasan et al., 2016), *Ift88^fl^* [MGI:3710185] (Haycraft et al., 2007), *mTmG* [MGI:3716464] (Muzumdar et al., 2007), *Sstr3-Gfp* [MGI: 5524281] (O’Connor et al., 2013), *Arl13b-Fucci2a* [MGI:6193732, RRID:IMSR_EM:12168] (Ford et al., 2018), and *Ptch1^LacZ/+^* [MGI1857447] (Goodrich et al., 1997). Note that *Ift88*^Δ^ is the deletion allele resulting from germline deletion of the conditional *Ift88^fl^*allele. Genotyping was performed on DNA extracted from ear punch via PCR (cycle: 95°C 0.5min, 60°C 0.5min, 72°C 1min) using the primers listed (Genotyping Primers Table).

### Genotyping Primers Table

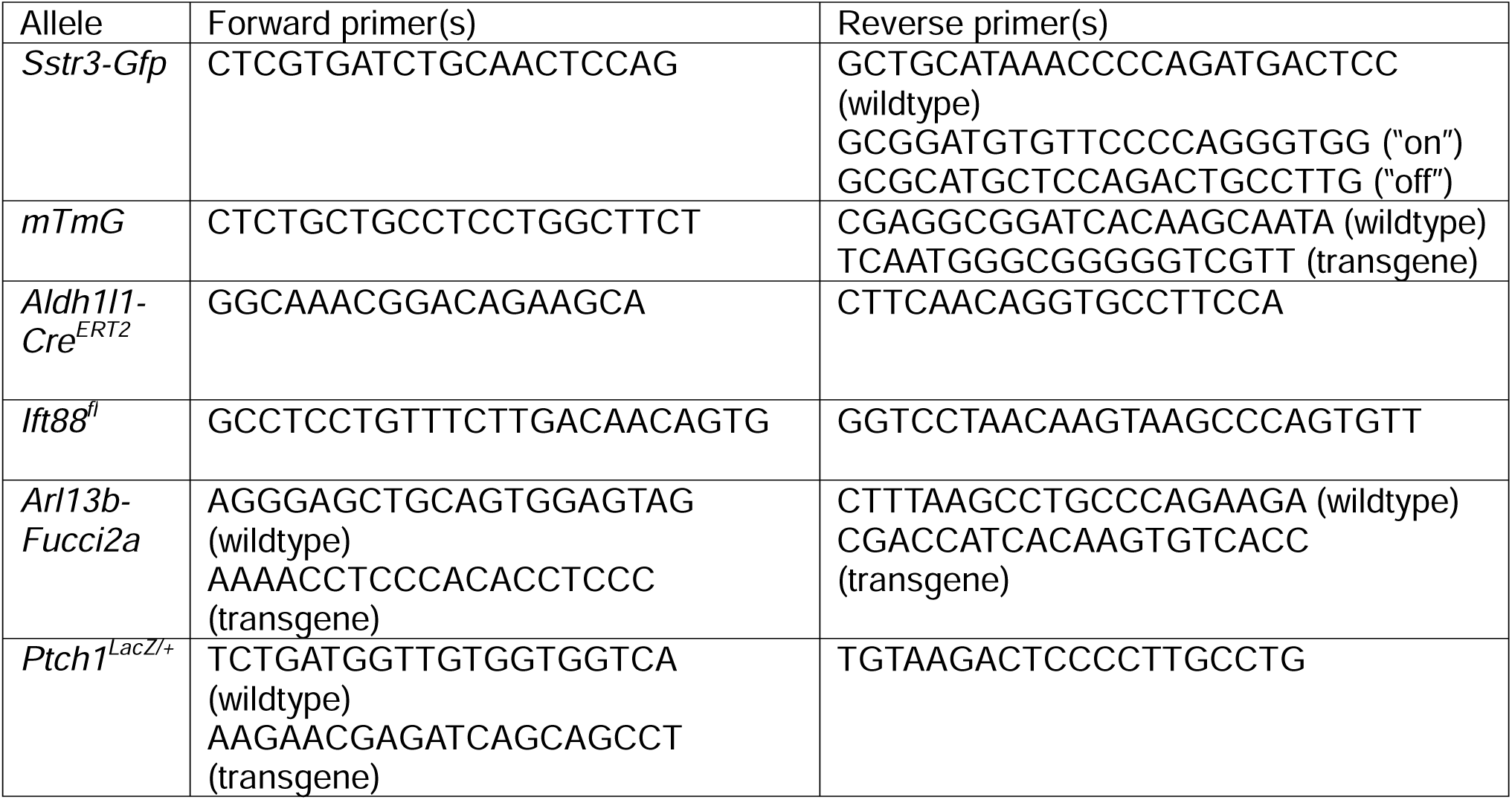

### Tamoxifen administration

Tamoxifen (Sigma T5648) stock solution was prepared once a month at a concentration of 10mg/ml in 100% EtOH and stored at -20°C. Each dose of tamoxifen was freshly prepared in corn oil the day of injection and dissolved using a speed vacuum centrifuge (Eppendorf Vacufuge Plus). To induce gene expression in immature astrocytes (*Ift88^Aldh-E15^*), 3mg tamoxifen/ 40g mouse weight in 300ul corn oil was administered intraperitoneally at E15.5 and E19.5 to the pregnant dam using a 1ml syringe and a 25G 5/8-inch needle (BD Biosciences) followed by subcutaneous administration of 3mg tamoxifen/ 40g mouse weight prepared in 20ul of corn oil at P0, P1, and P2 to pups using a 1/2ml syringe with attached 29G 1/2-inch needle (Covidien). To induce gene expression in mature astrocytes (*Ift88^Aldh-P0^*), 3mg tamoxifen/ 40g mouse weight in 300ul of corn oil was administered intraperitoneally at P0 and P1 to the dam using a 1ml syringe and a 25G 5/8-inch needle. To induce sparse gene expression in mature astrocytes (*Ift88^Aldh-P1^*), 1.25mg tamoxifen/ 40g mouse weight tamoxifen in 300ul corn oil was administered intraperitoneally at P1 to the dam using a 1ml syringe and a 25G 5/8-inch needle.

### Caesarean section and cross fostering

A Caesarean section was performed on tamoxifen-treated, timed-pregnant dams at E20.5. CD-1 mice were used as foster dams. Briefly, the experimental dams were euthanized via cervical dislocation and the pups were dissected, placed on a heating pad, and tapped gently until they could breathe on their own. Pups were transferred to a CD-1 foster dam and integrated with the foster dam’s litter.

### Tissue harvesting

P21 mice were euthanized by isoflurane inhalation followed by a trans-cardiac perfusion with ice-cold 1x phosphate-buffered saline (PBS) and ice-cold 4% paraformaldehyde (PFA). P2-P8 mice were euthanized by decapitation. Brains were harvested following perfusion or decapitation and drop-fixed in 4% PFA overnight. Fixed tissue was washed with 1x PBS and incubated with 30% sucrose in 0.1 M phosphate buffer overnight at 4C until the tissue sank.

Samples were washed in optimal cutting temperature (OCT) compound to remove sucrose (Tissue-Tek OCT, Sakura Finetek), embedded in OCT, frozen on dry ice, and stored at -20°C.

### EdU administration

EdU (Invitrogen A10044) stock solution was prepared at a concentration of 50mg/ml in sterilized 1x PBS. EdU solution was administered subcutaneously in a volume of 20ul to pups at P2 and P3 using a 1/2ml syringe with attached 29G 1/2-inch needle (Covidien). Tissue was processed the same as for immunofluorescent staining and EdU labeling was visualized using a homemade AlexaFluor-647 Click-IT EdU kit. Sections were washed with 1X Tris-Buffered Saline (TBS) to remove OCT, permeabilized with PBS-T (.3% Triton X-100), incubated in Click-It Reaction (1M Tris-HCl pH 8.5, 1M CuSO4, Alexa 647-Azide 1:1000 Invitrogen A10277, ddH20, and 0.5M sodium ascorbate) protected from light, washed in 1X TBS, and then followed by immunofluorescent staining.

### Immunofluorescent staining (IF)

OCT-embedded tissues were sectioned at 40µm using a cryostat microtome and placed directly on microscope slides (Fisherbrand Superfrost). Sections were let to air-dry and either processed immediately or stored at -20°C. Sections stored at -20°C were brought to room temperature before starting IF. Sections were first rehydrated in 1X TBS, permeabilized in 1% SDS, and blocked in antibody wash (1% heat inactivated goat serum, 0.1% Triton X-100 in 1X TBS).

Sections were incubated overnight at 4°C with primary antibodies (Primary Antibody Table). Sections were washed with cold antibody wash and incubated for one hour at room temperature in the dark with secondary antibodies (goat anti-chicken AlexaFluor 488; goat anti-chicken AlexaFluor 633; goat anti-mouse AlexaFluor 555 (IgG2a); goat anti-rabbit AlexaFluor 647; Hoechst 33342; all 1:500 dilution, ThermoFisher). Sections were washed with cold antibody wash and then mounted with a glass coverslip using ProLong Gold (ThermoFisher) mounting media. Slides cured overnight at room temperature in the dark and were stored short term at 4°C or long term at -20°C. Slides were imaged on a BioTek Lionheart FX automated microscope and Nikon A1R confocal microscope.

### Primary Antibody Table

**Primary Antibody Table:**
| **Antibody** | **Dilution** | **Company** | **Catalog #** |
| --- | --- | --- | --- |
| mouse anti-ARL13B | 1:500 | Neuromab | N295B/66 |
| chicken anti-AC3 | 1:500 | Encor | CPCA-ACIII |
| rabbit anti-SOX9 | 1:500 | Millipore | AB5535 |
| chicken anti-GFP | 1:8,000 | Abcam | ab13970 |
| rabbit anti-KI67 | 1:250 | Abcam | ab16667 |
| rabbit anti-RFP | 1:500 | Rockland | 600-401-379 |
| rabbit anti-GFAP | 1:500 | Abcam | ab7260 |
| chicken anti-β-Gal | 1:2000 | Abcam | ab9361 |

### Fluorescent *in-situ* hybridization assays

Mice were euthanized at the indicated age by cervical dislocation and brain tissue was harvested and immediately frozen in optimal cutting temperature compound and stored at - 80°C. Tissue was sectioned at 18µm using a cryostat microtome on microscope slides (Fisherbrand Superfrost) and immediately stored at -80°C. The Multiplexed HCR RNA-FISH assay was used according to manufacturer’s instructions for mouse fresh frozen tissue (Molecular Instruments). Briefly, slides were fixed in 4% PFA , dehydrated in a series of ethanol washes and incubated with experimental probes in a humidified chamber at 37°C overnight. The following customs probes were created to bind all known mRNA variants of *Aldh1l1* (B1), *Gli1*-37 probe sets and *Gli1*-20 probe sets (B2), *Ptch1* (B3). Slides were washed and then incubated in hairpin amplifiers (B1-488, B2-647, B3-555) for 4 hours in a humidified dark chamber at room temperature. Slides were washed and then mounted with a glass coverslip using Prolong Diamond Antifade mount (Thermo Fisher), stored in the dark at 4°C overnight, and imaged after 24 hours using a Nikon A1R confocal microscope.

### Quantification of astrocyte area, volume, and branching

Z-stack images of brain sections expressing *Aldh1l1-Cre^ERT2^;mTmG* were captured at 0.5um intervals using a Nikon A1R confocal microscope. Analysis of astrocyte morphology was performed using IMARIS surface rendering.

### Quantification of fluorescent in-situ hybridization

Analysis of all fluorescent *in-situ* images was performed using the HALO FISH module (Indica Labs). HALO generated counts of individua puncta (single transcript) for each segmented nuclei. Cells with greater than 10 transcripts for *Aldh1l1* were identified as astrocytes, and the average number of transcripts expressed in astrocytes for each target was calculated.

### Quantification of cilia

Z-stack images of brain sections stained for ciliary markers were captured at 2um intervals. Analysis of cilia was performed on maximum projection images using cell counting in Fiji/ImageJ.

### FACS purification of mouse cortical astrocytes

Astrocytes were purified following the protocol as published (Zhang et al., 2016). Briefly, the cortex was dissected from mice at ages P8 and P21 and dissociated with papain (Worthington Biochemical, LS003126, 20U/ml) in enzyme solution at 34°C with 5% C0_2_ for 45 minutes or 60 minutes, respectively. Then, the tissue was washed with a low inhibitor solution and triturated to dissociate cells. High inhibitor solution was added to the cell suspension and centrifuged for 5 minutes at 500g. The cell pellet was resuspended in FACS buffer (0.2% FBS in 1X PBS w/o Ca+ or Mg+) and passed through a 40um filter. Cell sorting was performed using the BD FACS Aria II (BD Biosciences) and GFP-positive and TdTomato-negative cells (approximately 100,000-200,000 cells/sample) were collected directly in 1mL of TRIzol-LS (Invitrogen, 10296010) and frozen at -80°C.

### RNA isolation and RNA sequencing analysis

Total RNA from sorted GFP-positive astrocyte was extracted using the miRNeasy Mini Kit (Qiagen, 217004) under the protocols of the manufacturer. The quality of RNA was determined using the Agilent Bioanalyzer Pico Chips and samples with RNA integrity number higher than 8 were used for sequencing. RNA samples were sent to Admera Health for library construction (SMARTseq V4 + Nextera XT) and sequencing (Illumina 2x 150bp reads, 40M read depth).

RNAseq reads were analyzed using the Galaxy web-platform where we assessed quality control using FASTQC, trimmed using Cutadapt, and mapped to the mouse genome version 10 (mm10). Read counts were generated using featureCounts and differential gene expression analysis was performed with edgeR. Genes with a fold change ≥ 0.75 (upregulated) or ≤ -0.75 (downregulated) and p-value ≤ 0.05 were considered differentially expressed. The functional enrichment analysis was performed using g:Profiler with g:SCS multiple testing correction and p-value < 0.05.

## RESULTS

### Primary cilia are present in a sub-population of developing astrocytes and predominantly express ARL13B

To characterize the properties of developing astrocyte cilia, we quantified the frequency and protein composition of astrocyte cilia in the prefrontal cortex. To label astrocyte cilia, we used *Aldh1l1-Cre^ERT2^*, a tamoxifen inducible Cre under an astrocyte-specific promoter, and the Cre-dependent cilia reporter *Sstr3-Gfp*. We administered tamoxifen embryonically to induce SSTR3-GFP expression in astrocytes (see methods and Figure 1 for details). This model permits detailed investigation of primary cilia specifically in the astrocyte lineage (Figure 1a).

**Figure 1:**
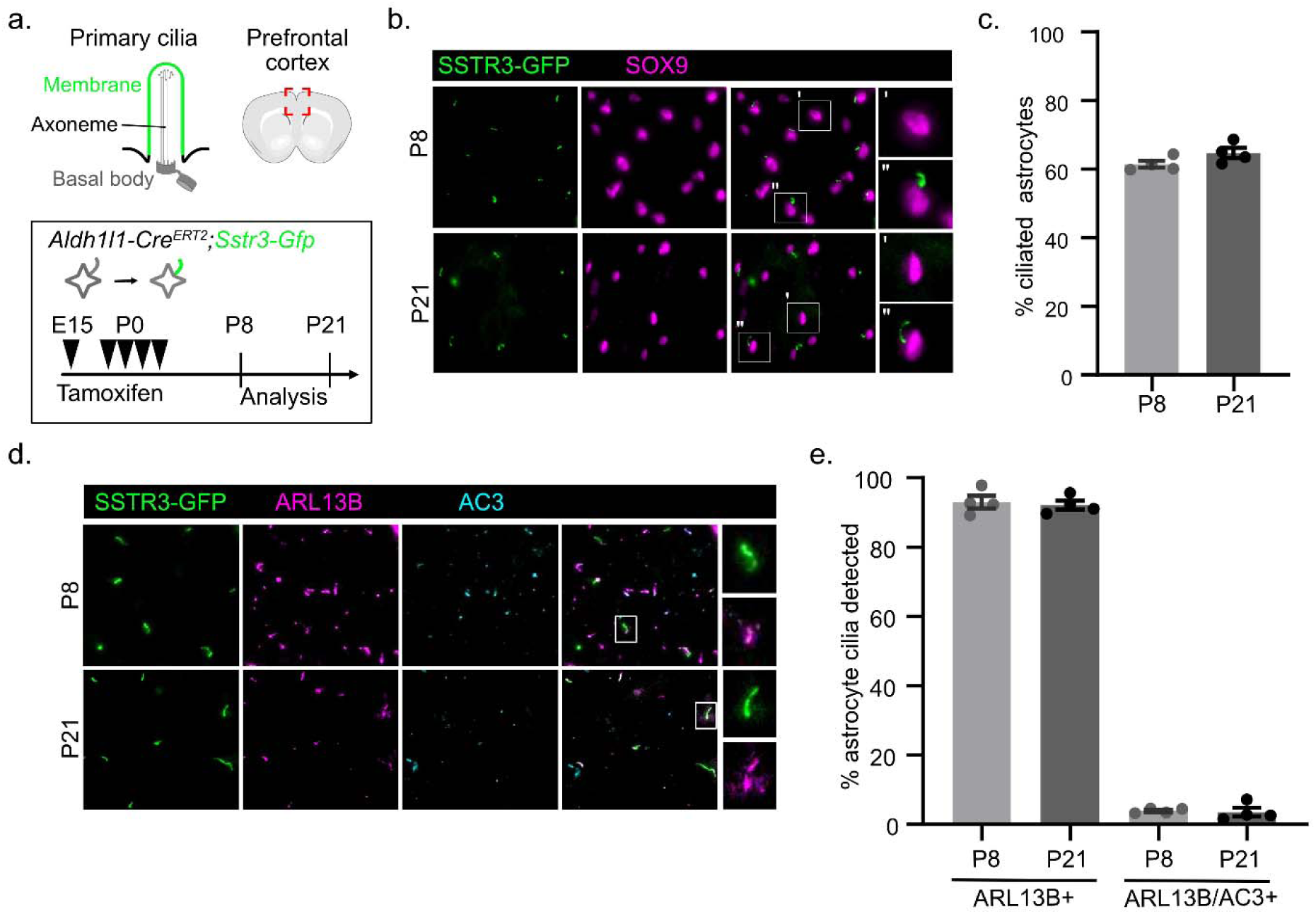
A subset of immature and mature astrocytes have cilia and predominantly express ARL13B. **(a)** Schematic of primary cilia, the prefrontal cortex (red, region of interest), and genetic labeling approach. **(b)** Representative image of GFP staining to detect SSTR3-GFP-positive astrocyte cilia and SOX9 staining to label astrocytes. **(c)** Quantification of the percent ciliated SOX9-positive astrocytes at P8 and P21. **(d)** Representative image of GFP staining to detect SSTR3-GFP-positive astrocyte cilia and ARL13B and AC3 staining at P8 and P21. **(e)** Quantification of the percent colocalization of ARL13B or ARL13B and AC3 (ARL13B/AC3) with SSTR3-GFP-positive astrocyte cilia. *n* = 4 animals. Data are expressed as the mean ± SEM.

We focused our analysis of cilia in P8 immature and P21 mature astrocytes. Using immunofluorescence staining, we quantified the number of astrocytes that possess GFP-positive cilia and the expression of ciliary proteins ARL13B and AC3 within astrocyte cilia (Figure 1b,d). We observed that 61.4% of immature astrocytes and 64.7% of mature astrocytes possess primary cilia (Figure 1c). This result indicates both that many developing astrocytes possess primary cilia and that about 1/3 of astrocytes lack them. In immature astrocytes, we observed that 93% of cilia express ARL13B and 4% of cilia express both ARL13B and AC3. In mature astrocytes, we observed that 92% of astrocyte cilia express ARL13B and 3.5% of cilia express both AC3 and ARL13B. There were no astrocyte cilia at either timepoint that expressed AC3 alone (Figure 1e). The astrocyte ciliation frequency and ciliary protein composition remain constant from P8 to P21 indicating astrocyte cilia are stable. These data show that a sub-population of developing astrocytes possess cilia and suggests that ARL13B is a reasonable marker for astrocyte cilia.

### Transcriptomic analysis of immature *Ift88^Aldh-E15^* astrocytes reveals changes in Shh signaling

The presence of cilia in developing astrocytes prompts whether cilia contribute specific functions to astrocyte biology. To determine the function of cilia during early stages of astrocyte development, we conditionally deleted the Intraflagellar transport 88 (*Ift88*) gene in the astrocyte lineage by generating *Aldh1l1-Cre^ERT2^*;*Ift88^f/^*^Δ^ mice. As IFT88 is required for cilia formation and maintenance of cilia structure, this strategy genetically ablates cilia in all Cre-expressing cells (Pazour et al., 2000). IFT88 is a stable protein so its deletion takes time to affect cilia; therefore, we used mice that already contain one deletion allele (*Ift88^f/^*^Δ^). To induce complete loss of cilia in immature astrocytes, we administered tamoxifen embryonically-referred to hereafter as *Ift88^Aldh-E15^*(see methods and Supp. Figure 1a for details). We determined when cilia are lost in the astrocyte lineage by incorporating the Cre-dependent cilia and cell cycle reporter *Arl13b-Fucci2a* (*AF2a*) into control and *Ift88^Aldh-E15^* mice. In the prefrontal cortex at P4, we observed 24% fewer astrocyte cilia and by P8 we found 93.8% fewer astrocyte cilia in *Ift88^Aldh-E15^;AF2a* compared to *AF2a* control mice (Supp. Figure 1a-c). These data illustrate that embryonic ablation of astrocyte cilia is gradual during the first postnatal week and that P8 is the earliest feasible timepoint to study the consequences of cilia ablation in immature astrocytes.

To examine the impact of loss of cilia on immature astrocyte gene expression, we isolated cortical astrocytes from P8 control and *Ift88^Aldh-E15^*mice and performed bulk RNA sequencing. We incorporated the reporter allele *mTmG,* which expresses membrane bound green fluorescent protein (GFP) upon Cre recombination. To obtain a pure population of astrocytes, we dissected the cortex, dissociated cells, and used FACS to isolate GFP-positive astrocytes. We observed 33 significantly downregulated genes and 47 significantly upregulated genes in *Ift88^Aldh-E15^* astrocytes using a Log FC cutoff of 0.75 and p-value < 0.05 (Figure 2a). The short differentially expressed gene list indicates that we captured specific changes in astrocytes likely to be a direct consequence of cilia ablation (Supp. Table 1). There were no biological processes identified when we performed gene ontology analysis for the upregulated genes.

**Figure 2:**
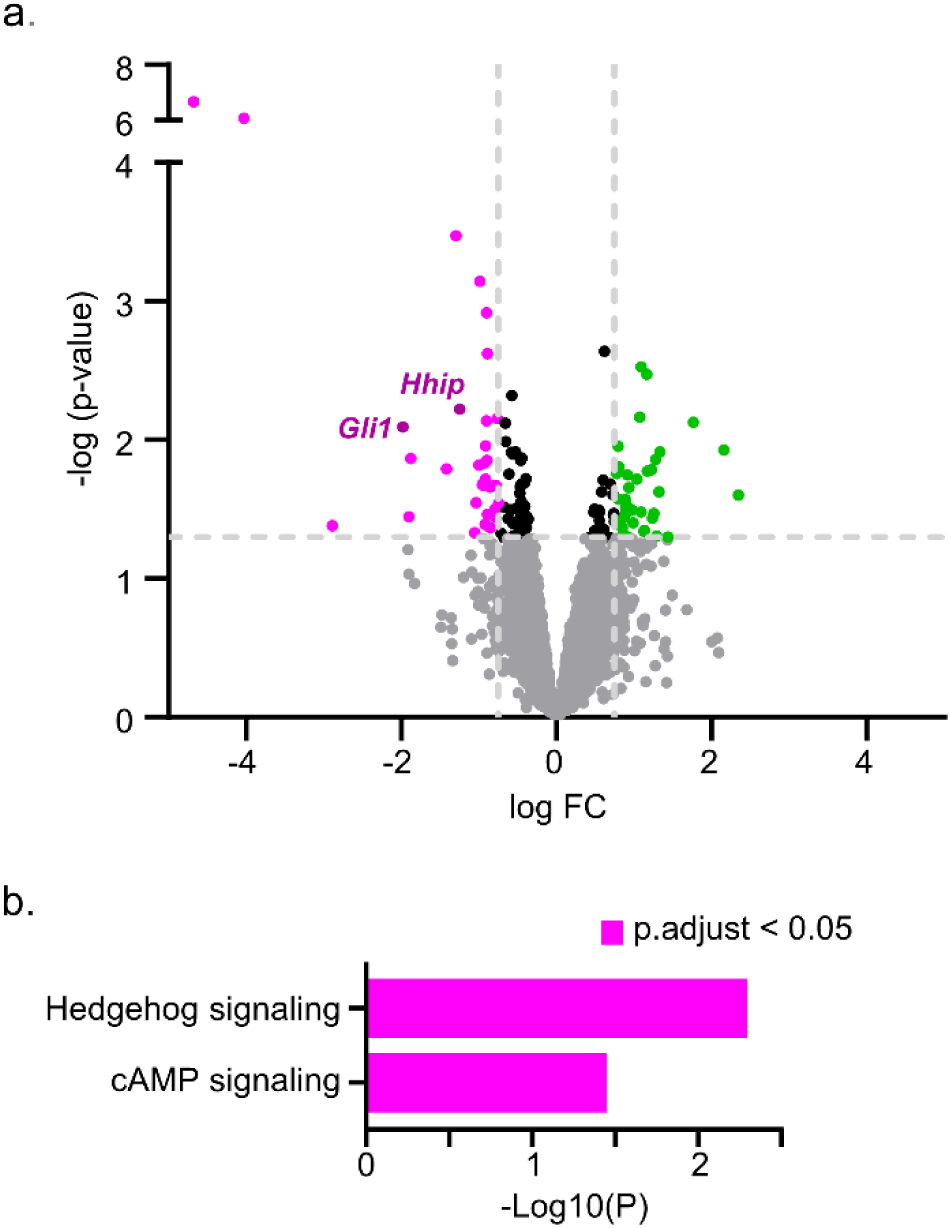
RNAseq analysis reveals changes in ciliary signaling in Ift88^Aldh-E15^. **(a)** Volcano plot showing differentially expressed genes in *Ift88^Aldh-E15^* versus control astrocytes, upregulated genes (green) and downregulated genes (magenta), log FC cutoff of 0.75 and p-value < 0.05. *n* = 5 animals. **(b)** Gene ontology terms enriched in downregulated genes.

Among the downregulated genes, we found enrichment for gene ontology terms related to cAMP signaling (*Atp2b4*) and Shh signaling (*Gli1, Hhip*) (Figure 2b). This finding indicates that IFT88 is required to regulate Shh signaling in immature astrocytes.

### Loss of cilia in immature astrocytes results in decreased Shh signaling *in vivo*

To gain spatial information about the transcriptomic changes, we used fluorescent *in situ* hybridization to investigate the Shh response in immature astrocytes at P8. We labeled astrocytes with the *Aldh1l1* probe and monitored Shh target genes with *Gli1* and *Ptch1* probes (Figure 3a). We observed an overall 70.4% decrease in *Gli1* transcript and 16.3% decrease in *Ptch1* transcript in astrocytes of *Ift88^Aldh-E15^* mice compared to control (Figure 3c,e). When we evaluated transcripts by the spatial location of astrocytes in the prefrontal cortex (PFC), we found that the *Gli1* transcript was significantly decreased in all layers, whereas *Ptch1* transcript downregulation was specific to layers I-IV (Figure 3b,d). The decrease in Shh target gene expression suggests that Shh activity is reduced in immature astrocytes upon IFT88 deletion.

**Figure 3:**
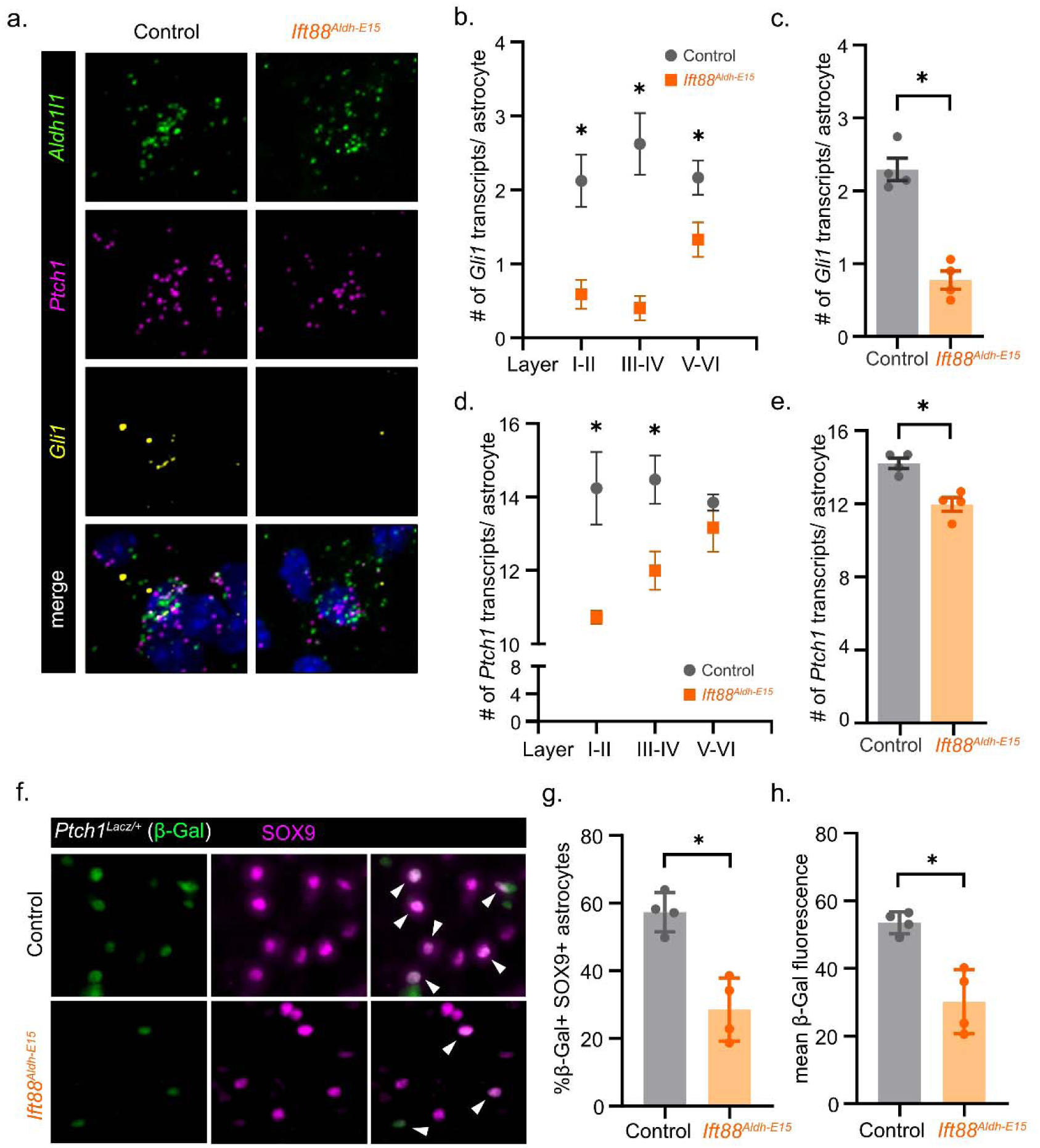
Ift88^Aldh-E15^ astrocytes display decreased Shh target gene expression and reduced Shh activity. **(a)** Representative image of fluorescent *in situ* hybridization staining for the astrocyte maker *Aldh1l1* and Shh target genes *Ptch1* and *Gli1* in control and *Ift88^Aldh-E15^* astrocytes. **(b)** Quantification of the average number of *Gli1* transcripts per astrocyte by PFC layer and **(c)** across all PFC layers in control and *Ift88^Aldh-E15^*. **(d)** Quantification of the average number of *Ptch1* transcripts per astrocyte by PFC layer and **(e)** across all PFC layers in control and *Ift88^Aldh-E15^*. **(f)** Representative image of β-Gal staining (green) to detect *Ptch1^LacZ/+^* and SOX9 staining (magenta) to label astrocytes in *Ptch1^LacZ/+^* control and *Ift88^Aldh-E15^; Ptch1^LacZ/+^* mice. **(g)** Quantification of the percent β-Gal-positive SOX9-positive astrocytes and **(h)** β-Gal mean fluorescence intensity in *Ptch1^LacZ/+^* control and *Ift88^Aldh-E15^; Ptch1^LacZ/+^* mice. *n* = 4 animals. Data are expressed as the mean ± SEM. Statistical analysis using *t*-test, *p-value < 0.05.

**Figure 4:**
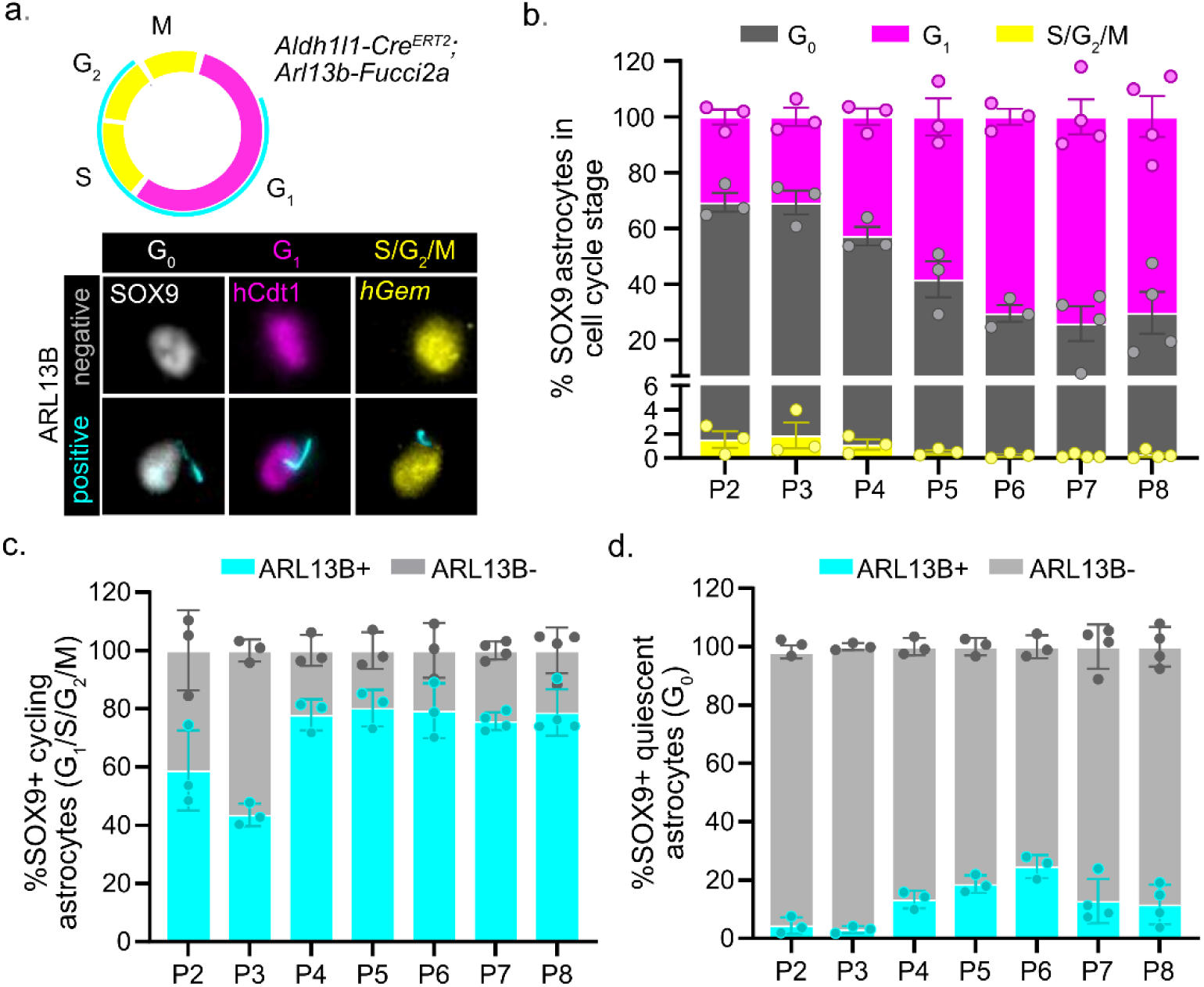
The relationship between cilia and the cell cycle during astrocyte proliferation. **(a)** Schematic of the *Arl13b-Fucci2a* reporter and representative image of astrocytes with or without cilia in each cell cycle stage. **(b)** Quantification of the percent of astrocytes in each cell cycle stage from P2-P8 in control pups. **(c)** Quantification of the percent of cycling astrocytes with or without cilia. **(d)** Quantification of the percent of quiescent astrocytes with or without cilia. *n* = 3-4 animals. Data are expressed as the mean ± SEM.

To test this directly, we used the *Ptch1^LacZ/+^* reporter allele, which expresses β-Galactosidase (β-Gal), as a readout for Shh activity. Using immunofluorescence staining, we quantified β-Gal expression in the PFC at P8 (Figure 3f). We observed a 50.2% decrease in the number of β-Gal-positive astrocytes and a 43.6% decrease in β-Gal fluorescence intensity in *Ift88^Aldh-E15^; Ptch1^LacZ/+^* compared to *Ptch1^LacZ/+^*control mice (Figure 3 g,h). Next, we determined whether the population of Shh-active cells corresponds to ciliated astrocytes. In *Ptch1^LacZ/+^* control mice, we observed that 94% of β-Gal-positive astrocytes possess cilia while only 33.4% of β-Gal-negative astrocytes possess cilia (Supp. Figure 2a). This suggests that the transcriptomic changes in Shh target genes are restricted to the sub-population of ciliated astrocytes. Taken together, these data establish that there is a significant reduction of Shh activity in *Ift88^Aldh-E15^* astrocytes and proposes that cilia mediate Shh activity in early postnatal astrocyte development.

### Cilia-cell cycle biosensor shows cilia are increased in cycling astrocytes compared to quiescent astrocytes

To examine the relationship between astrocytes and cilia during proliferation, we incorporated the cilia-cell cycle biosensor *Arl13b-Fucci2a* (hereafter called *AF2a*) into *Aldh1l1-Cre^ERT2^* mice. The *AF2a* reporter expresses a Cre-dependent tricistronic cassette containing *Arl13b-Cerulean*, *hCdt1-mCherry*, and *hGem-mVenus* which labels cilia, G_1_ cells, and S/G_2_/M cells, respectively (Figure 4a). To induce AF2a expression in astrocytes, we administered tamoxifen embryonically (*AF2a^Aldh-E15^*). Using immunofluorescence staining, we quantified the number of SOX9-positive astrocytes in each stage of the cell cycle from P2-P8 in the PFC. The absence of the cell cycle markers and presence of SOX9 labels astrocytes in G_0_. In control *AF2a^Aldh-E15^*, we found that the greatest number of S/G_2_/M astrocytes are present from P2-P4 (2.42%) compared to P5-P8 (0.32%). From P2-P8, we observed a decrease in the number of G_0_ astrocytes (64.5% at P2 to 29.3% at P8) and a proportionate increase in the number of G_1_ astrocytes (34.9% at P2 to 70.4% at P8) (Figure 4b). These results are consistent with the peak window of astrocyte proliferation (Ge et al., 2012). Within the cycling astrocyte population, an average of 70.9% of astrocytes express cilia from P2-P8 (Figure 4c). In contrast, an average of 4.9% of astrocytes in the quiescent population express cilia from P2-P8 (Figure 4d). This finding indicates that cilia are present in proliferating astrocytes and suggests that cilia could serve important roles during astrocyte proliferation but not quiescence.

### Loss of cilia in immature astrocytes results in decreased astrocyte proliferation

We next explored whether cilia play a role in astrocyte proliferation. To determine the function of cilia during astrocyte proliferation, we quantified the number of dividing astrocytes in *Ift88^Aldh-E15^* and control mice. We incorporated the *mTmG* reporter allele to label the astrocyte lineage and administered EdU at P2 and P3 to label dividing cells and their progeny. Using immunofluorescence staining, we quantified the number of EdU-positive or Ki67-positive astrocytes in the PFC at P4. We observed 26.6% fewer EdU-positive astrocytes and 18.8% fewer Ki67- positive astrocytes in *Ift88^Aldh-E15^;mTmG* compared to *mTmG* control mice (Figure 5a,b). To examine whether the population of astrocytes was changed as a result of decreased proliferation, we quantified the number of astrocytes in defined areas of the PFC (spatial density) from P4-P8. We found no changes in the spatial density of astrocytes in *Ift88^Aldh-E15^* compared to control mice (Supp. Figure 2b).

**Figure 5:**
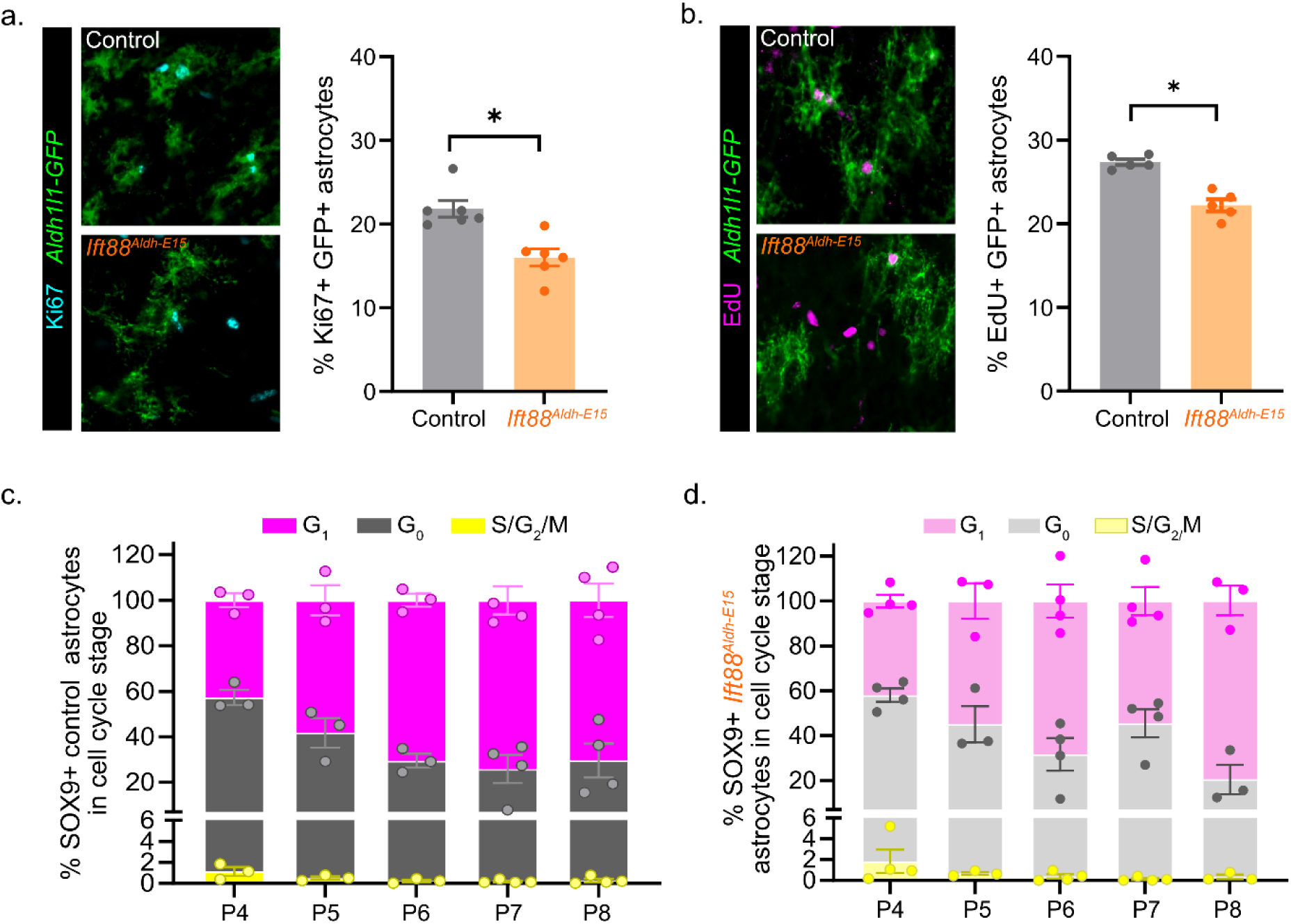
Ift88^Aldh-E15^ astrocytes display decreased proliferation. **(a)** Representative image of staining for Ki67 and GFP to label proliferating astrocytes and quantification of the percent Ki67-positive astrocytes in P4 *mTmG* control *Ift88^Aldh-E15^;mTmG*. **(b)** Representative image of staining for EdU and GFP to label proliferating astrocytes and quantification of the percent EdU-positive astrocytes in P4 *mTmG* control and *Ift88^Aldh-E15^;mTmG*. **(c)** Quantification of the percent of astrocytes in G_0_, G_1_, or S/G_2_/M from P4-P8 in *AF2a* control. **(d)** Quantification of the percent of astrocytes in G_0_, G_1_, or S/G_2_/M from P4-P8 in *Ift88^Aldh-E15^;AF2a*. *n* = 4-6 animals. Data are expressed as the mean ± SEM. Statistical analysis using *t*-test. *p-value < 0.05.

A potential explanation for the diminished proliferation in astrocytes lacking IFT88 is that the cells stall in G_1_. To analyze astrocyte cell cycle progression, we incorporated the *Arl13b-Fucci2a* (*AF2a)* reporter allele to label astrocyte cell cycle stage. We used immunofluorescence staining to quantify the number of astrocytes in each stage of the cell cycle in *Ift88^Aldh-E15^*;*AF2a* and *AF2a* control pups from P4-P8. We observed no difference in the number of astrocytes in each stage of the cell cycle at any timepoint (Figure 5c,d). This argues that the loss of IFT88 does not disrupt astrocyte cell cycle progression, despite cilia being required during astrocyte proliferation.

### Transcriptomic analysis of mature *Ift88^Aldh-P0^* astrocytes reveals changes in developmental signaling pathways

To investigate what role cilia play in maturing astrocytes, we shifted the timing of *Ift88* deletion in our astrocyte cilia genetic ablation model. To target maturing astrocytes, we administered tamoxifen postnatally in *Aldh1l1-Cre^ERT2^;Ift88^f/^*^Δ^ mice-referred to hereafter as *Ift88^Aldh-P0^*(see methods and Supp. Figure 1d for details). We determined when cilia are lost in the astrocyte lineage by incorporating the *AF2a* reporter into control and *Ift88^Aldh-P0^* mice. At P10, we observed 80% fewer astrocyte cilia and by P14 we found 95% fewer astrocyte cilia in *Ift88^Aldh-P0^;AF2a* compared to *AF2a* control mice (Supp. Figure 1d-f). This model results in loss of astrocyte cilia by the second postnatal week of astrocyte development allowing us to specifically evaluate cilia function during astrocyte maturation at P21.

To determine the function of cilia during astrocyte maturation, we used the same transcriptomic approach as in the immature astrocyte model. Briefly, we used FACS to isolate GFP-positive astrocytes from the cortex of P21 *Ift88^Aldh-P0^;mTmG* and *mTmG* control mice and performed bulk RNA sequencing. We observed 107 significantly downregulated genes and 15 significantly upregulated genes in *Ift88^Aldh-P0^* astrocytes using a Log FC cutoff of 0.75 and p-value ≤ 0.05 (Figure 6a). We performed gene ontology analysis and found that the downregulated genes are enriched in broad biological processes including regulation of signal transduction, regulation of cell communication, developmental processes, and anatomical structure (Figure 6b). There were no biological processes identified in the upregulated genes when we performed gene ontology analysis. Several downregulated genes are related to specific developmental signaling pathways associated with cilia including Shh (*Gli1* and *Hhip*), Wnt (*Axin1, Dact3, Znrf3*), and TGFβ/BMP (*Smad4, Insm1,* and *Rcor1*) (Corbit et al., 2008; Huangfu et al., 2003; Lancaster et al., 2011; Mönnich et al., 2018) (Supp. Table 1). These data uncover several cilia-linked signaling pathways that may contribute to astrocyte maturation. This result further suggests that IFT88 loss impacts Shh signaling in astrocytes and motivates investigation of Shh activity in mature astrocytes.

**Figure 6:**
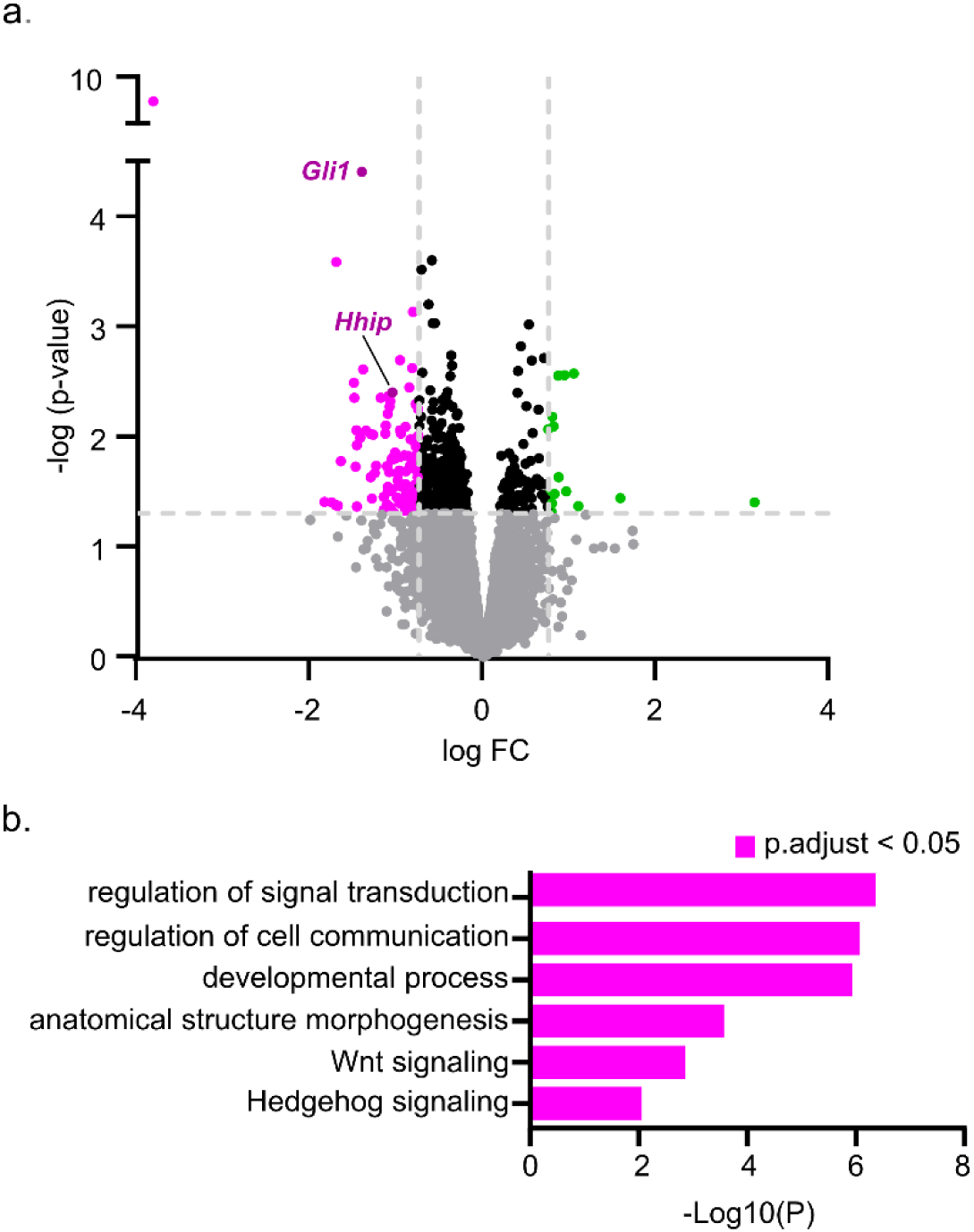
RNAseq analysis reveals changes in developmental signaling pathways in Ift88^Aldh-P0^. **(a)** Volcano plot showing differentially expressed genes in *Ift88^Aldh-P0^* versus control astrocytes, upregulated genes (green) and downregulated genes (magenta), log FC cutoff of 0.75 and p-value < 0.05. **(b)** Gene ontology terms enriched in downregulated genes.

### Loss of mature astrocyte cilia causes decreased Shh signaling

To obtain spatial information about the transcriptomic changes, we used fluorescent *in situ* hybridization to examine the Shh response in mature astrocytes at P21. We labeled astrocytes with the *Aldh1l1* probe and monitored Shh target genes with *Gli1* and *Ptch1* probes. We observed a 50.4% decrease in *Gli1* transcripts and 17.8% decrease in *Ptch1* transcripts specifically in PFC layers V-VI in *Ift88^Aldh-E15^* compared to control astrocytes (Figure 7a-e). These results illustrate that astrocytes in deeper cortical layers are responsible for the decrease in Shh target genes following IFT88 deletion.

**Figure 7:**
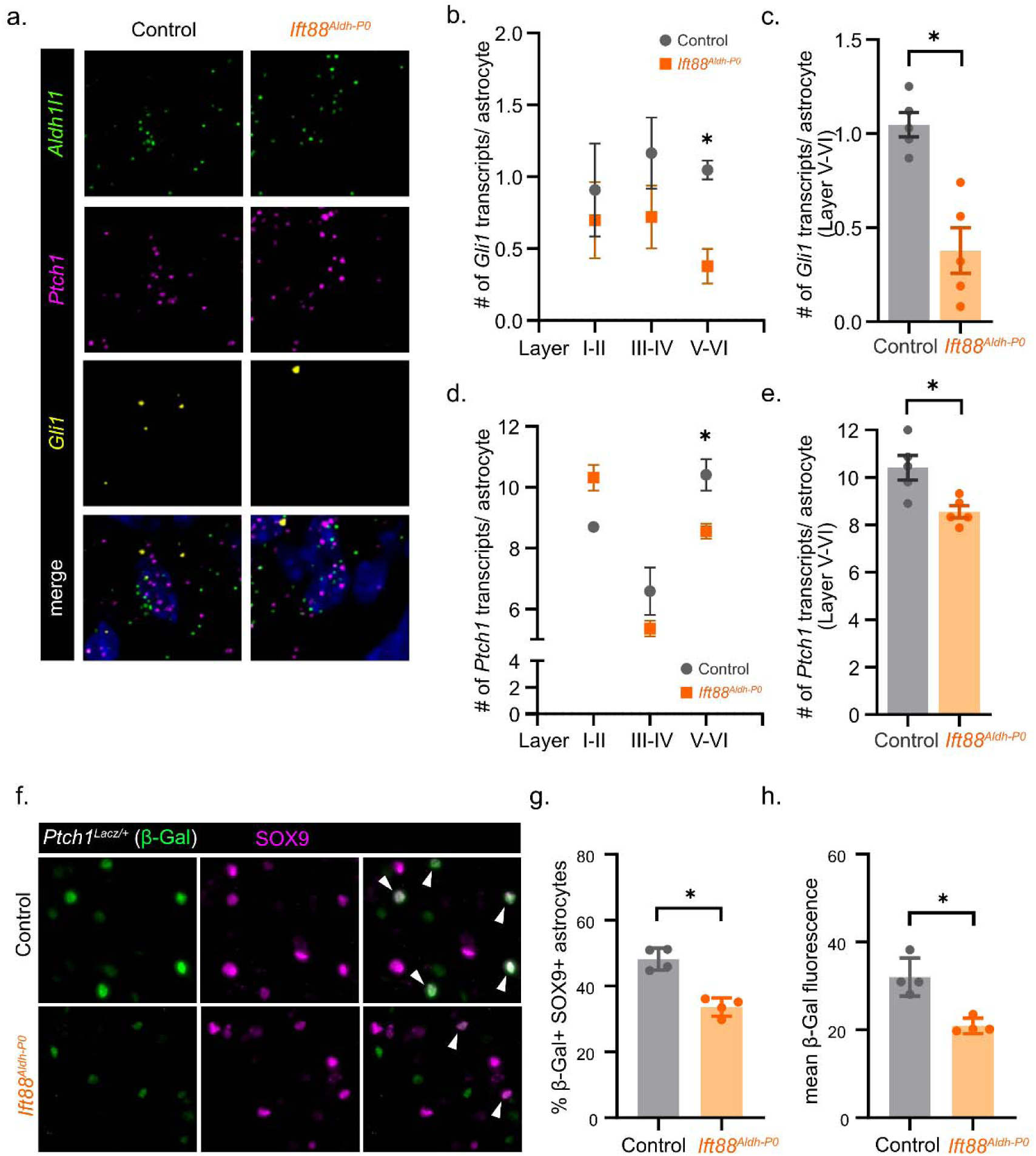
I**ft88^Aldh-P0^ deep PFC layer astrocytes display decreased Shh target gene expression and reduced Shh activity. (a)** Representative image of fluorescent *in situ* hybridization staining for the astrocyte maker *Aldh1l1* and Shh target genes *Ptch1* and *Gli1* in PFC layer V-VI control and *Ift88^Aldh-P0^* astrocytes. **(b)** Quantification of the average number of *Gli1* transcripts per astrocyte by PFC layer and **(c)** PFC layer V-VI in control and *Ift88^Aldh-P0^*. **(d)** Quantification of the number of *Ptch1* transcripts per astrocyte in each PFC layer and **(e)** PFC layer V-VI in control and *Ift88^Aldh-P0^*. **(f)** Representative image of β-Gal staining (green) to detect *Ptch1^LacZ/+^* and SOX9 staining (magenta) to label astrocytes in *Ptch1^LacZ/+^* control and *Ift88^Aldh-P0^; Ptch1^LacZ/+^* mice. **(g)** Quantification of the percent β-Gal-positive SOX9- positive astrocytes and **(h)** β-Gal mean fluorescence intensity in *Ptch1^LacZ/+^* control and *Ift88^Aldh-P0^; Ptch1^LacZ/+^* mice. *n* = 4 animals. Data are expressed as the mean ± SEM. Statistical analysis using *t*-test, *p-value < 0.05.

The decreased expression of Shh transcriptional target genes predicts that Shh activity is reduced in mature astrocytes lacking IFT88. To test this *in vivo*, we again incorporated the *Ptch1^LacZ/+^* reporter and used immunofluorescence staining to quantify β-Gal expression at P21 in the PFC (Figure 7f). We observed 30.2% fewer β-Gal-positive astrocytes and a 39.5% decrease in β-Gal fluorescence intensity in *Ift88^Aldh-P0^*;*Ptch1^LacZ/+^* compared to *Ptch1^LacZ/+^* control mice (Figure 7g,h). This finding confirms that loss of IFT88 results in reduced Shh activity.

Interestingly, we observed no difference in the number of β-Gal-positive astrocytes or β-Gal fluorescence intensity by cortical layer (Supp. Figure 3a). Furthermore, in *Ptch1^LacZ/+^* control mice, we found that 88.6% of β-Gal-positive astrocytes possess cilia while only 24.8% of β-Gal-negative astrocytes are ciliated (Supp. Figure 3b). Together, these data support that astrocyte cilia regulate Shh activity and that cilia likely function in mature astrocytes.

### Loss of primary cilia in mature astrocytes causes enlarged astrocyte processes

To determine the function of cilia during astrocyte maturation, we evaluated astrocyte morphology. We incorporated the *mTmG* reporter allele to label the membrane of astrocyte processes. To visualize individual astrocytes, we administered a low dose of tamoxifen at P1 to achieve sparse labeling (hereafter referred to as *Ift88^Aldh-P1^*). We found that this dose is sufficient to induce loss of cilia while labeling individual astrocytes (Supp. Figure 4). Using immunofluorescence staining, we performed morphological analysis of P21 astrocytes in the PFC (Figure 8a,b). We observed a 23% increase in the membrane area and 22% increase in membrane volume of *Ift88^Aldh-P1^*;*mTmG* compared to *mTmG* control astrocytes (Figure 8c,d).

**Figure 8:**
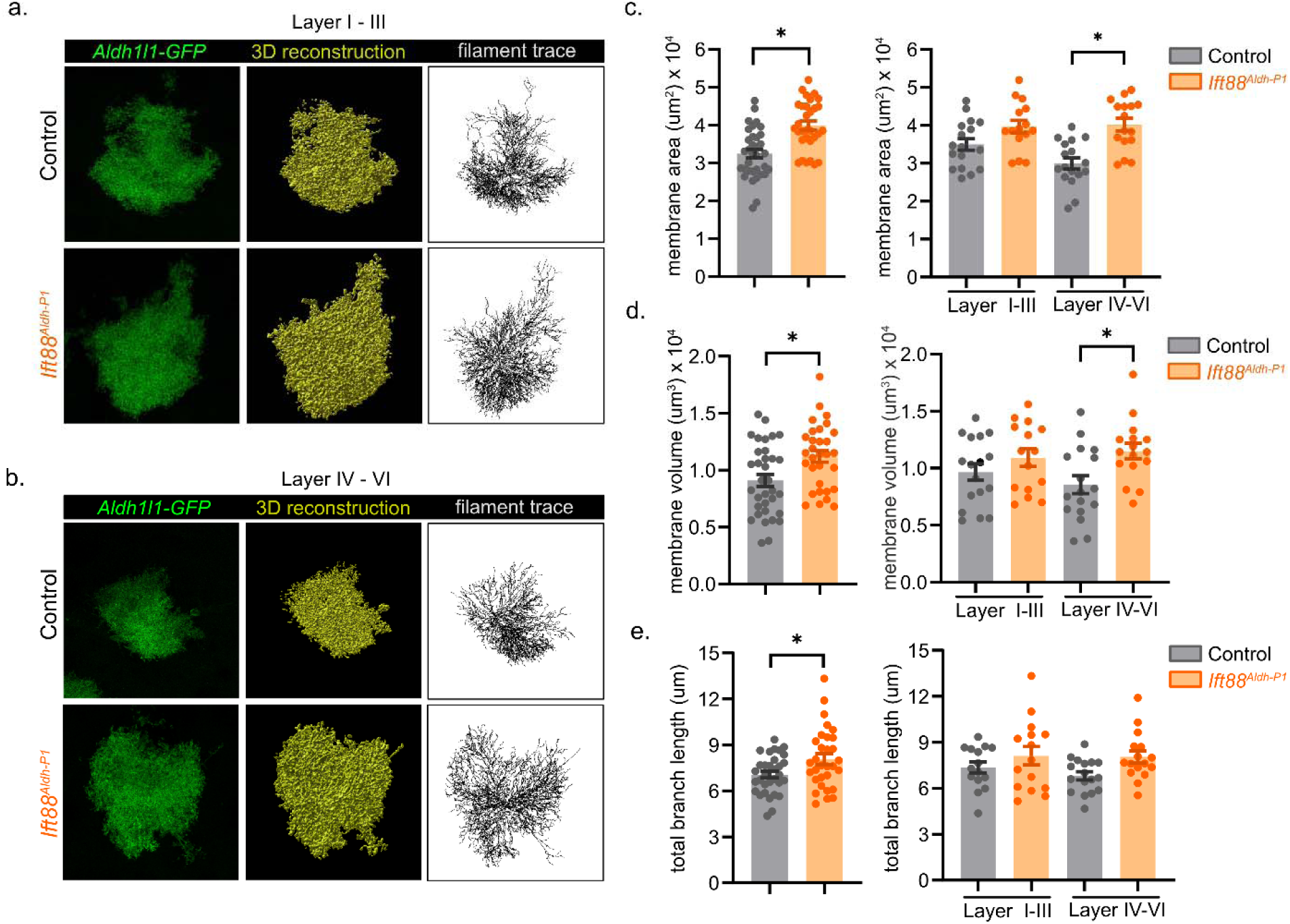
Ift88^Aldh-P1^ astrocytes display increased membrane and branch structure in deep cortical layers. **(a,b)** Representative confocal images of *Aldh1l1-GFP* astrocytes, 3-D reconstruction, and filament trace from PFC layer I-III and layer IV-VI in P21 control and *Ift88^Aldh-P1^* astrocytes. **(c)** Quantification of membrane area and **(d)** membrane volume in total astrocytes and by PFC layer in control and *Ift88^Aldh-P1^* astrocytes. **(e)** Quantification of total branch length in total astrocytes and by PFC layer in control and *Ift88^Aldh-P1^* astrocytes. *n* = 3 animals, 30 astrocytes per genotype (15 each for layer I-III and layer IV-VI). Data are expressed as the mean ± SEM. Statistical analysis using *t*-test or one-way ANOVA followed by Turkey’s test for multiple comparisons, *p-value < 0.05.

*Ift88^Aldh-P1^* astrocytes located in layers IV-VI exhibited a significantly larger membrane area and volume compared to *Ift88^Aldh-P1^* astrocytes in layers I-III (Figure 8c,d). Furthermore, we observed a 14.3% increase in total branch length of *Ift88^Aldh-P1^;mTmG* compared to *mTmG* control astrocytes (Figure 8e). However, the total branch length was not altered in a cortical layer-specific manner. These data show that astrocyte processes increase upon IFT88 deletion and that these changes are more pronounced in astrocytes located in deep cortical layers. Together, these findings indicate that cilia play a role in coordinating astrocyte morphology.

## DISCUSSION

Here, we applied genetic tools to visualize and manipulate cilia in developing astrocytes. We showed that a subset of developing astrocytes in the prefrontal cortex possess cilia. We genetically ablated cilia in immature and mature astrocytes and found that Shh activity was decreased, indicating that astrocytes require cilia for Shh signaling throughout their development. In immature astrocytes, we showed that proliferating astrocytes correspond to the ciliated population and loss of astrocyte cilia results in decreased proliferation. In mature astrocytes, we found that loss of cilia causes enlarged astrocyte processes implicating cilia function in astrocyte maturation. Our work uncovers novel roles for cilia in developing astrocytes and broadens our understanding of the molecular mechanisms underlying astrocyte development.

Previous studies of astrocyte cilia focused on defining their presence in the brain as the functions of astrocyte cilia were unknown (Kasahara et al., 2014; Sipos et al., 2018; Sterpka & Chen, 2018). Here, we extended these findings to define astrocyte cilia in early stages of postnatal development in the PFC. Using two genetic labeling approaches, we demonstrated that a subset of astrocytes possess cilia at P8 and P21. This was surprising given that the majority of astrocytes have a cilia in the primary visual cortex at P36 (Ott et al., 2023).

Furthermore, in adult mice, almost all cortical astrocytes are ciliated (Wang et al., 2024). It is possible that the presence of cilia in astrocytes varies across different brain regions and at different timepoints. Our finding relied on the expression of fluorescently tagged cilia-localized proteins to label astrocyte cilia and will need to be corroborated by electron microscopy.

Nonetheless, our work prompts interesting questions about whether cilia confer specific functions within the astrocyte population.

During postnatal astrocyte proliferation, the ciliated astrocyte population corresponds to cycling astrocytes. This raises the possibility that the population of astrocytes undergoing local proliferation build cilia and then retain cilia after proliferation is complete. It is currently unclear how this sub-population of ciliated astrocytes is established and how this population changes at adult stages. Our work reveals previously unknown biology about developing astrocyte cilia and highlights the unique relationship between astrocyte cilia and the cell cycle.

ARL13B is one of the few ciliary proteins that we have tools to detect *in vivo* in the brain. ARL13B is a small atypical GTPase that plays roles in cilia structure, protein trafficking, and Shh signaling (Caspary et al., 2007). While ARL13B is expressed in astrocyte cilia, little is understood about its function in developing astrocytes (Kasahara et al., 2014; Sipos et al., 2018; Sterpka & Chen, 2018). In adult mice, ARL13B functions in astrocytes to maintain their morphology and regulate neural circuit function (Wang et al., 2024). We found that ARL13B is the predominant ciliary protein expressed in developing astrocytes. We observed that very few astrocyte cilia express AC3, which is consistent with AC3 expression in neuronal cilia (Bishop et al., 2007). This work is important as the field currently lacks an astrocyte marker. In addition to defining ARL13B function during astrocyte development, other outstanding questions include what additional signaling proteins localize to astrocyte cilia and what functions they serve.

Shh is a notable developmental signaling pathway that serves critical roles in many stages of neuron development (Baudoin et al., 2012; Falcón-Urrutia et al., 2015; Ferent et al., 2019; Harwell et al., 2012; Huangfu et al., 2003; Wechsler-Reya & Scott, 1999). Importantly, Shh is active in astrocyte progenitors and continues to function in a subpopulation of adult astrocytes (Gingrich et al., 2022). In both models of cilia ablation (*Ift88^Aldh-E15^ and Ift88^Aldh-P0^*), we observed changes in Shh transcriptional targets. These models of astrocyte cilia ablation use *Ift88* deletion as a proxy for cilia loss. *Ift88* mutants have a well-established loss of cilia phenotype in many tissues (Boldizar et al., 2024; Huangfu et al., 2003; Kutomi et al., 2022; Murcia et al., 2000; Pant et al., n.d., 2024; Tian et al., 2017; Tong et al., 2014). Here, we showed that Shh target genes are decreased *in vivo* and that Shh activity is reduced upon loss of astrocyte cilia. These changes in Shh activity are driven by the ciliated astrocytes, which comprise the majority of Shh-active astrocytes. We observed that the ciliated and Shh-responsive astrocytes are dispersed throughout all PFC layers. Some Shh response remained in *Ift88^Aldh-E15^* and *Ift88^Aldh-P0^* astrocytes which could reflect that Shh activity perdures following loss of cilia. Alternatively, there could be de-repression of Shh target genes in astrocytes that never possessed cilia, which has been observed in other tissues (Lex et al., 2022). Our work demonstrates the requirement of cilia for Shh activity in developing astrocytes *in vivo*. Recent work showed that impaired ciliary signaling in later stages of astrocyte development causes Shh-dependent changes in astrocyte gene expression (Wang et al., 2024). Together, these findings establish that when astrocyte cilia are present, they function to transduce Shh signaling.

Cilia function in cellular proliferation by regulating cell cycle progression and transducing proliferative signals such as Shh (Chizhikov et al., 2007; Jonassen et al., 2008; Yeh et al., 2013). We observed reduced proliferation in *Ift88^Aldh-E15^*astrocytes revealing a link between cilia and astrocyte proliferation. Despite the decreased astrocyte proliferation at P4, we did not observe changes in the spatial density of astrocytes in the PFC. While there is some loss of astrocyte cilia at P4, many astrocytes still have cilia. Therefore, the remaining ciliated astrocytes could compensate to establish proper numbers of astrocytes. We predicted that Ioss of cilia would cause astrocytes to stall in G_1_, however we did not observe changes in cell cycle progression. It is possible that our approach could not capture the precise kinetics of cell cycle progression. We identified Shh as an important ciliary signal in immature astrocytes, which suggests Shh could serve as a proliferative cue in astrocytes. *N-myc* is a proliferative transcriptional target of Shh signaling; however, this gene was not differentially expressed in *Ift88^Aldh-E15^* astrocytes (Oliver et al., 2003). Future studies are needed to uncover how cilia regulate astrocyte proliferation.

In addition to Shh signaling, we found changes in TGFβ/BMP and Wnt signaling in *Ift88^Aldh-P0^*astrocytes. In fact, TGFβ/BMP and Wnt signaling play a role in regulating astrocyte morphology (Scholze et al., 2014; Szewczyk et al., 2023). However, the relationships between cilia and TGFβ/BMP or Wnt signaling are complex and often tissue specific. Wnt pathway components localize to cilia where Wnt signaling is modulated through different spatial compartments and protein interactions (Corbit et al., 2008; Gerdes et al., 2007; Lancaster et al., 2011). In some tissues, loss of cilia results in an expansion of Wnt signaling, while in other tissues, loss of cilia has no impact on Wnt signaling (Corbit et al., 2008; Huang & Schier, 2009; Ocbina et al., 2009; Simons et al., 2005). Cilia localize TGFβ/BMP receptors on their membrane and regulate the balance of TGFβ/BMP signaling through receptor internalization at the ciliary base (Anvarian et al., 2019; Clement et al., 2013; Mönnich et al., 2018). Additionally, these pathways interact, potentially obscuring the role cilia play in each. Shh and Wnt form antagonistic reciprocal interactions, TGFβ/BMP modulates Shh signaling, and Wnt and TGFβ/BMP synergistically regulate transcriptional targets (Guo & Wang, 2009; Kumar et al., 2021; Rios et al., 2004). There could also be cilia-independent functions of *Ift88* in modulating these signaling pathways. Our work reveals a novel connection between Wnt/TGFβ/BMP and astrocyte cilia that merits further investigation.

Shh signaling influences astrocyte morphology (Hill et al., 2019; Wang et al., 2024; Xie et al., 2022). Importantly, these studies demonstrated that the timing of Shh activity and balance of pathway output differentially impacts astrocyte morphology When we examined astrocyte morphology, we found that *Ift88^Aldh-P1^* astrocytes have enlarged processes specifically in deep layer PFC astrocytes. Our work expands on the role of Shh signaling in astrocyte maturation by showing the requirement of cilia. Furthermore, in *Ift88^Aldh-P0^* astrocytes, we found differences in the expression of Shh target genes specifically in deep PFC layers. This is in agreement with the finding that Shh ligand is enriched in layer V neurons (Harwell et al., 2012). Other work demonstrated that Shh signaling regulates astrocyte gene expression in a layer-dependent manner and disruption of Shh signaling in astrocytes impacts the morphology of deep cortical layer astrocytes (Wang et al., 2024; Xie et al., 2022). Together, these data indicate that Shh signaling influences astrocyte maturation in a spatial-dependent manner.

Astrocytes communicate among themselves and with other cells during the development of the nervous system. Primary cilia serve as signaling centers that facilitate cellular communication. Our work shows that cilia are necessary to regulate signals that are important for developing astrocytes. The presence of cilia in sub-populations of astrocytes may reflect functional among astrocytes. Ciliary signaling is an important aspect of astrocyte communication and will be important to study in the context of nervous system dysfunction.

## Data availability statement

The data that support the findings of this study are openly available and have been submitted to the GenBank database under accession number PRJNA1165324.

## Acknowledgements

We are grateful to Caitlin Sojka and Alexia King for technical help in purifying astrocytes and their associated RNA. Additionally, we thank Quinn Eastman, Alyssa Long, Tiffany Terry, Hanh Truong, and Robert Van Sciver for comments on the manuscript.

