## Supplemental Figure for "Primary cilia shape postnatal astrocyte development through Sonic Hedgehog signaling"

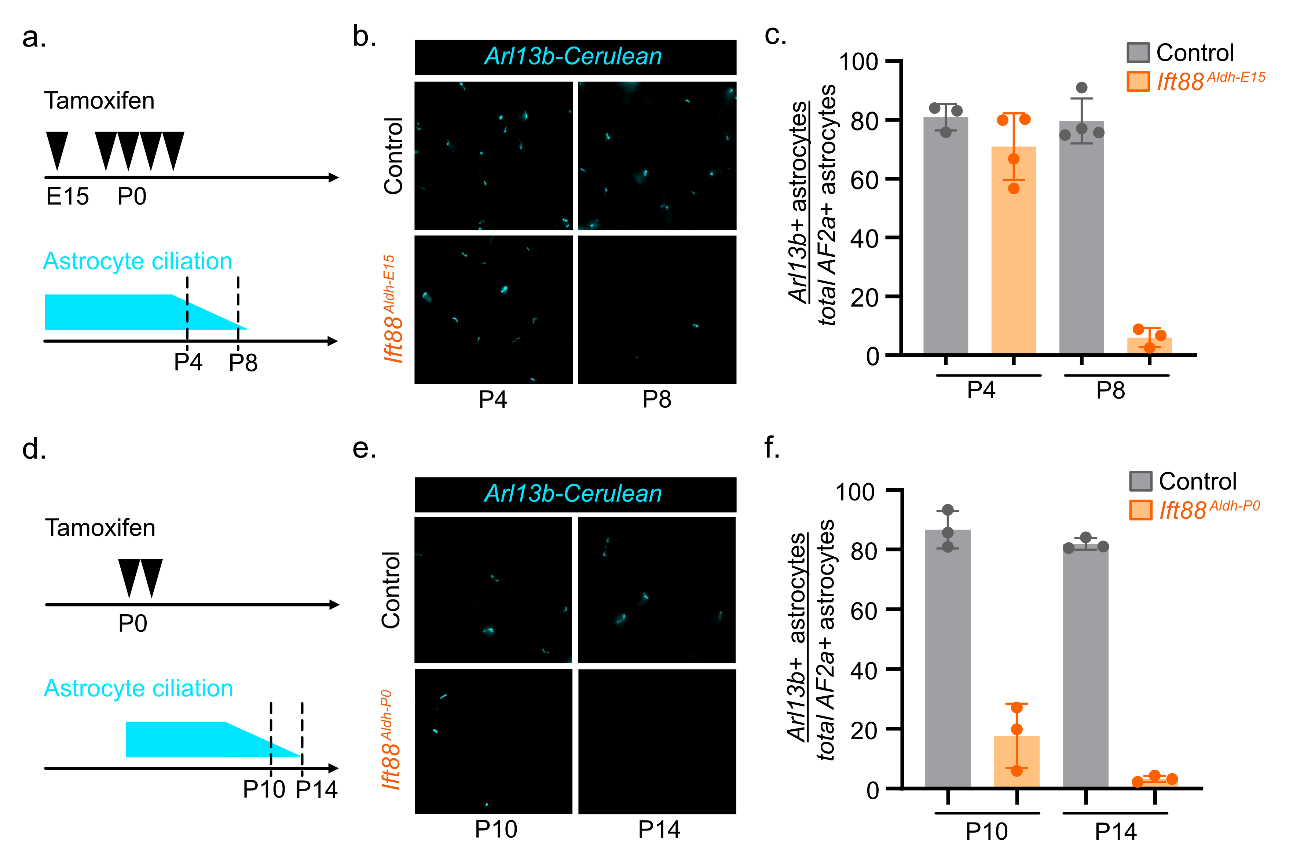

**Supplemental Figure 1: Kinetics of genetic cilia ablation models for immature and mature astrocytes (a)** Schematic of tamoxifen protocol for embryonic induction of cilia ablation to target immature astrocytes. **(b)** Representative image of Arl13b-cerulean (cilia reporter in AF2a) detection at P4 and P8 in AF2a control and Ift88^Aldh-E15^;AF2a mice. **(c)** Quantification of the percent Arl13b-positive over total AF2a- positive astrocytes at P4 and P8 in AF2a control and Ift88^Aldh-E15^;AF2a mice. **(d)** Schematic of tamoxifen protocol for postnatal induction of cilia ablation to target mature astrocytes. **(e)** Representative image of Arl13b-cerulean detection at P10 and P14 in AF2a control and Ift88^Aldh-P0^; AF2a mice. **(f)** Quantification of the percent Arl13b-positive over total AF2a- positive astrocytes at P10 and P14 in AF2a control and Ift88^Aldh-P0^; AF2a mice. n = 3-4 animals. Data are expressed as the mean ± SEM.

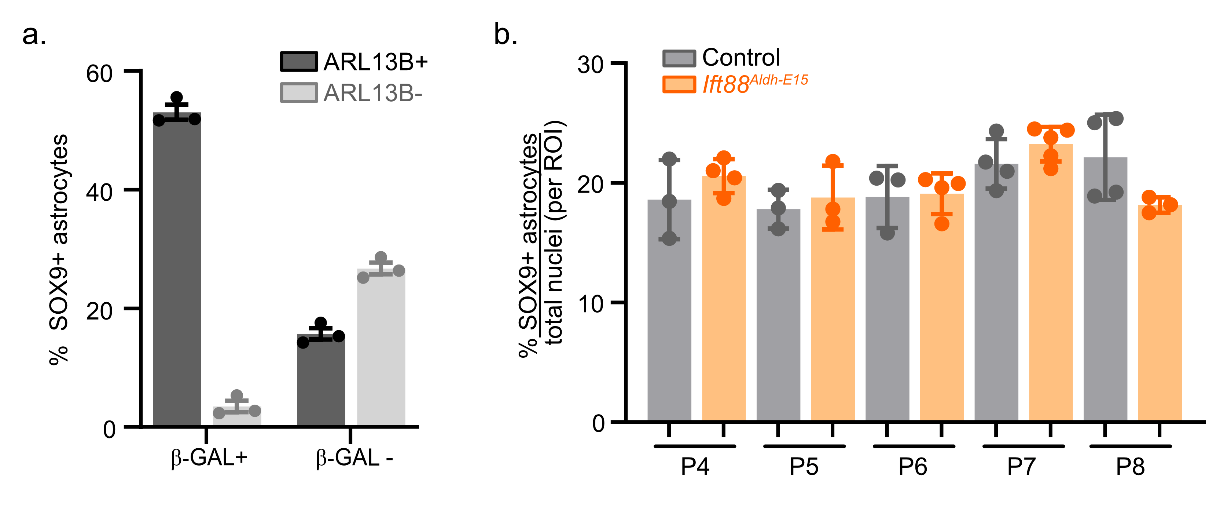

**Supplemental Figure 2: (a)** Quantification of the percent SOX9- positive astrocytes that are β-Gal- positive or β-Gal-negative and ARL13B- positive or ARL13B- negative in Ptch1^LacZ/+^ control mice at P8. **(b)** Quantification of the percent SOX9- positive astrocytes relative to the total number of nuclei detected in defined areas of the PFC (ROI) in control and Ift88^Aldh-E15^. n = 3-4 animals. Data are expressed as the mean ± SEM.

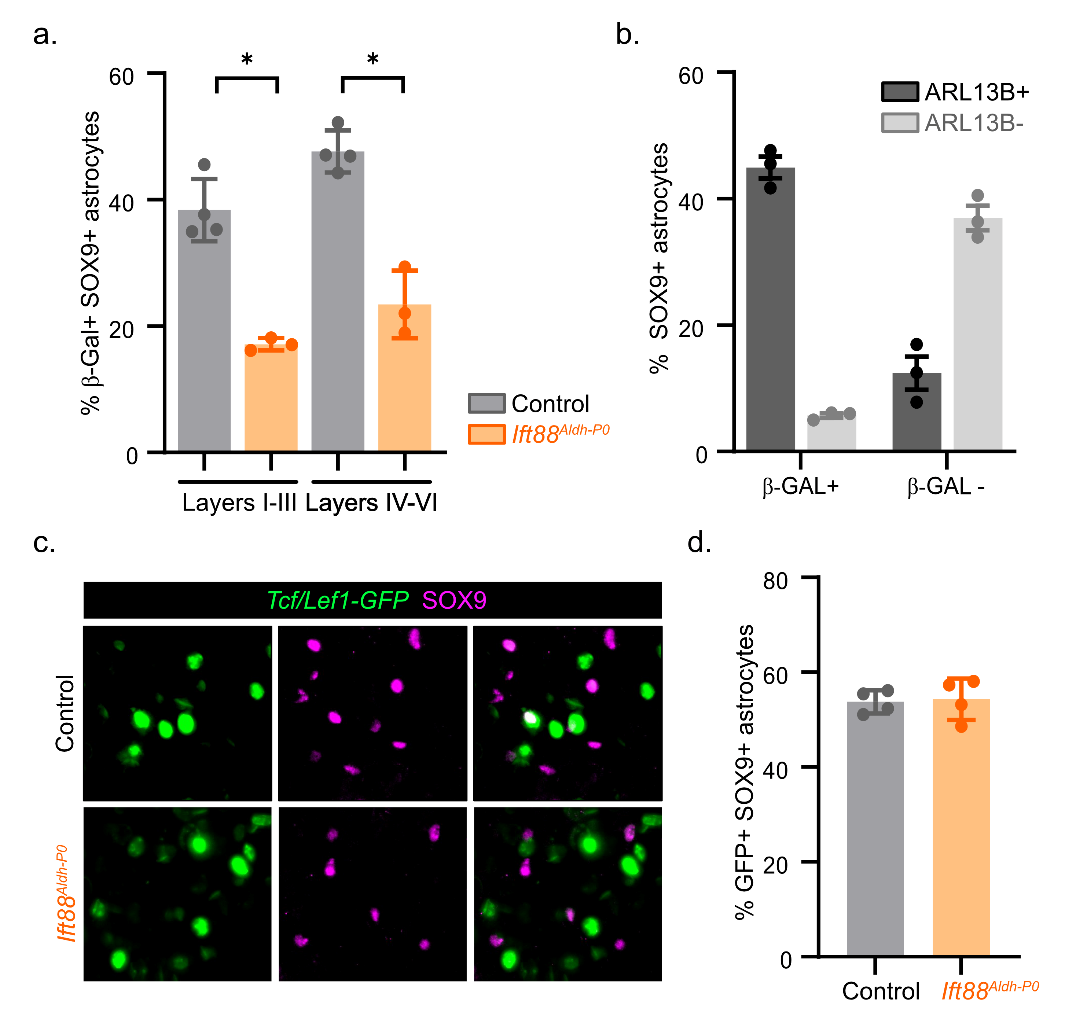

**Supplemental Figure 3: (a)** Quantification of the percent SOX9- positive and β-Gal- positive astrocytes in Layers I-III or Layers IV-VI in Ptch1^LacZ/+^ control and Ift88^Aldh-P0^;Ptch1^LacZ/+^ mice. **(b)** Quantification of the percent SOX9- positive astrocytes that are β-Gal- positive or β-Gal- negative and ARL13B- positive or ARL13B- negative in Ptch1^LacZ/+^ control mice at P21. n = 3-4 animals. Data are expressed as the mean ± SEM. Statistical analysis using t-test, p-value < 0.05.

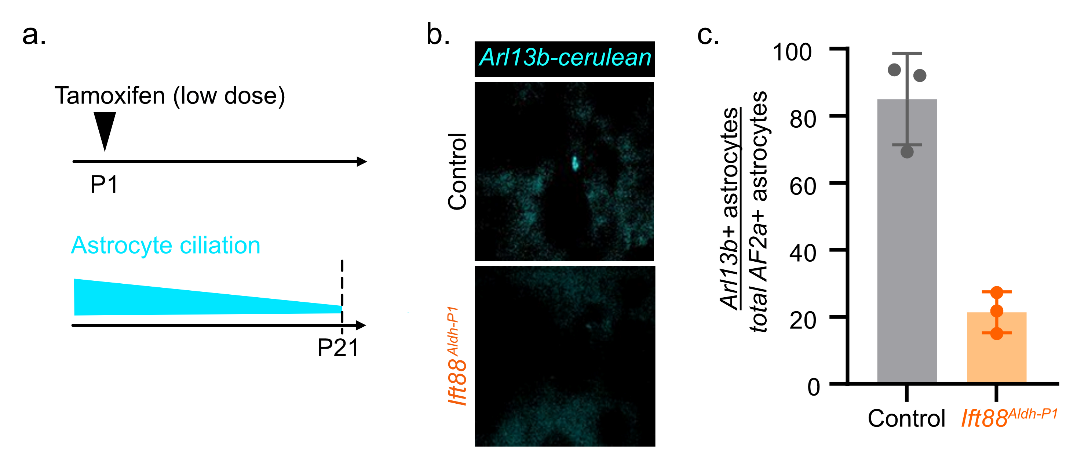

**Supplemental Figure 4: Validation of cilia loss in sparse labeling model of mature astrocytes (a)** Schematic of postnatal sparse labeling protocol. **(b)** Representative image of Arl13b-cerulean detection in AF2a control and Ift88^Aldh-P1^;AF2a mice. **(c)** Quantification of the percent Arl13b-positive over total AF2a- positive astrocytes at P21 in AF2a control and Ift88^Aldh-P1^;AF2a mice. n = 3 animals. Data are expressed as the mean ± SEM.

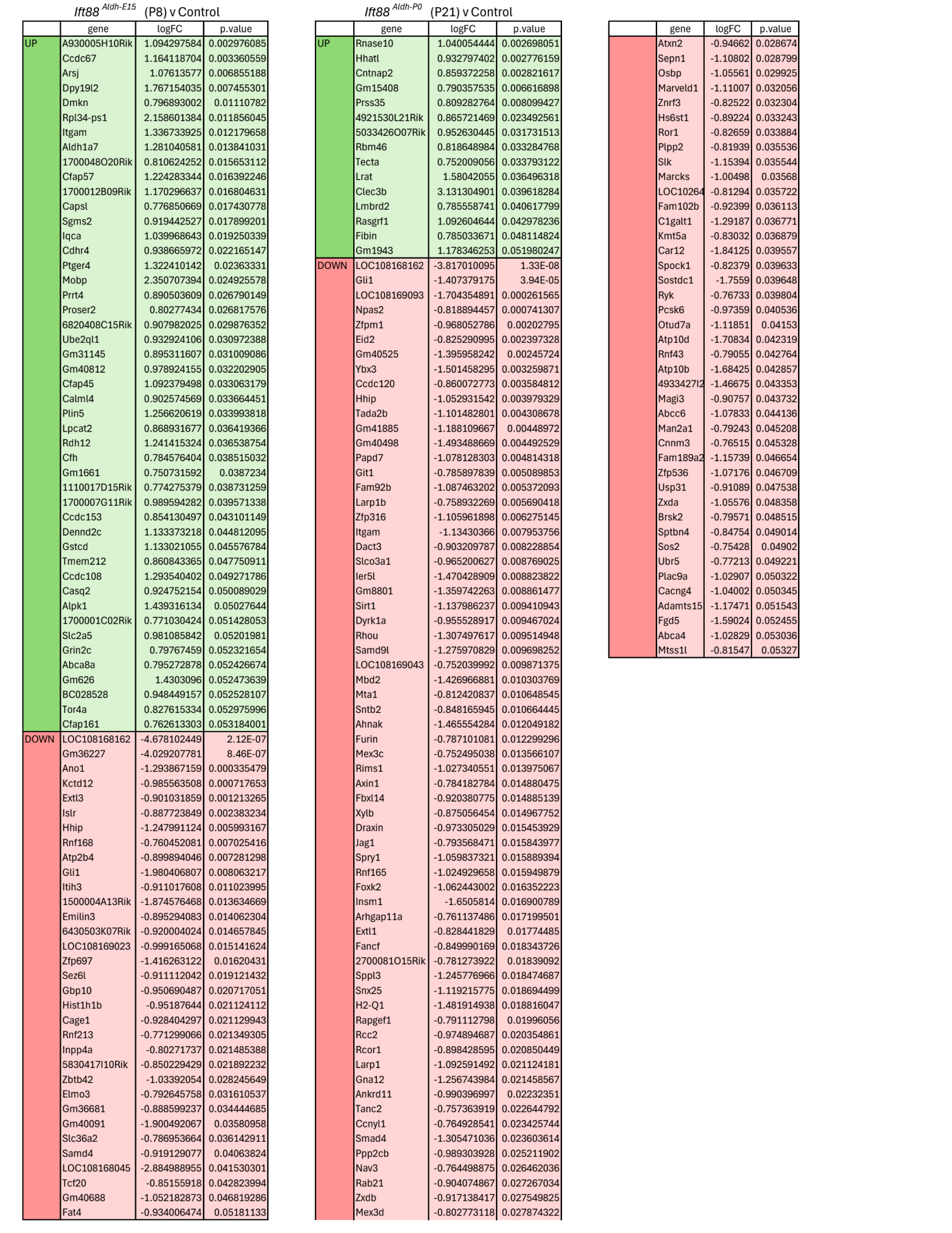

**Supplemental Table 2: Differentially expressed genes in loss of cilia astrocytes at P8 and P21**
